# Stretching the skin of the fingertip creates a perceptual and motor illusion of touching a harder spring

**DOI:** 10.1101/203604

**Authors:** Mor Farajian, Raz Leib, Tomer Zaidenberg, Ferdinando Mussa-Ivaldi, Ilana Nisky

## Abstract

We investigated how artificial tactile feedback in the form of a skin-stretch affects perception of stiffness and grip force adjustment. During interactions with objects, information from kinesthetic and tactile sensors is used to estimate the forces acting on the limbs. These enable the perception of the mechanical properties of objects to form, and the creation of internal models to predict the consequences of interactions with these objects, such as feedforward grip-force adjustments to prevent slippage. Previous studies showed that an artificial stretch of the skin of the fingertips can produce a linear additive effect on stiffness perception, but it remains unclear how such stretch affects the control of grip force. Here, we used a robotic device and a custom-built skin-stretch device to manipulate kinesthetic and tactile information. Using a stiffness discrimination task, we found that adding artificial tactile feedback to a kinesthetic force can create the illusion of touching a harder spring which affects both perception and action. The magnitude of the illusion is linearly related to the amplitude of the applied stretch. We also isolated the contribution of tactile stimulation to the predictive and reactive components of grip force adjustment, and found that unlike in other cases of perceptual illusions, the predictive grip force is modulated consistently with the perceptual tactile-induced illusion. These results have major implications for the design of tactile interfaces across a variety of touch applications such as wearable haptic devices, teleoperations, robot-assisted surgery, and prosthetics.

**Significance Statement:** Sensing forces, using kinesthetic and tactile modalities, is important for assessing the mechanical properties of objects, and for acting the objects while stabilizing grasp against slippage. A major challenge in understanding the internal representations that allow for a predictive grip force control during contact with objects is to dissociate the contribution of tactile and kinesthetic stimuli. To date, this contribution was investigated only in impaired cases either through local anesthesia or in patients with sensory impairment. Our study demonstrates using a programmable mechatronic device that artificially applied skin-stretch creates an illusion of a greater load force that affects grip force control and stiffness perception. These results are applicable in tactile technologies for wearable haptic devices, teleoperation, robot-assisted surgery, and prosthetics.

## Introduction

During everyday interactions with objects, people concurrently control and sense position and force in perceptual tasks such as when assessing the stiffness of an object using a tool (LaMotte 2000), as well as in actions such as controlling tool manipulation and adjusting the grip force – the perpendicular force between the digits and the object. Perception and action are tightly coupled in the sensorimotor system. For example, perception of the mechanical properties of the environment is important in planning future actions. At the same time, natural haptic exploration of the environment is active in that we move and probe the environment to create haptic perception. Since we do not possess stiffness sensors, the perception of stiffness is based on the integration of position and force signals (Jones, Hunter 1993, Kuschel et al. 2010, Gurari, Kuchenbecker & Okamura 2013).

There are two major types of forces sensing modalities in the human body – kinesthetic and tactile. When holding a pen or scalpel, kinesthetic position and force information is generated by muscle spindles (length and shortening velocity of the muscle) and Golgi tendon organs (tension in the tendon). Tactile information is generated by cutaneous mechanoreceptors that respond to a deformation of the skin (stretch, vibration, pressure). Steady skin deformation is obtained from the slowly adapting mechanoreceptors, whereas the rapidly adapting mechanoreceptors sense transient motion on the skin and alert the brain in case of slippage (Kandel et al. 2000). People with impaired tactile sensibility suffer from the absence of important information that is needed to plan and control object manipulation, such as magnitude and direction of contact forces, shapes of contact surfaces, and friction (Johansson, Flanagan 2009). Understanding the integration of the kinesthetic and tactile modalities is also important in the development of technology that displays force information to users of robotic devices such as teleoperation (Nisky, Mussa-Ivaldi & Karniel 2013, Schorr et al. 2015), robot-assisted surgery (Quek et al. 2015b, Pacchierotti, Prattichizzo & Kuchenbecker 2015), and prosthetics (Akhtar et al. 2014).

Using anesthesia and passive stimulation via a robot, Srinivasan and Lamotte (Srinivasan, LaMotte 1996) identified the unique role of tactile information in the discrimination of subtle differences in softness. However, passive or anesthetized touch is different from active touch (Gwilliam et al. 2014). Recently, there has been a significant advancement in the development of devices for tactile stimulation of the finger pads (Prattichizzo, Pacchierotti & Rosati 2012, Quek et al. 2014). In most of these devices, a moving tactor (a pin with a flat high-friction top) is in contact with the skin and causes an artificial skin deformation. This deformation emulates the effects observed when a person’s extremities interact with real objects. These devices can augment perception without increasing the kinesthetic force, and thus have the potential to overcome some of the practical limitations of providing force feedback, such as closed-loop stability. They are also instrumental to investigating the combination of tactile and kinesthetic information in a variety of scenarios. For example, a study in which artificial skin-stretch was added to a kinesthetic force showed that tactor displacement-induced skin-stretch had a linear additive effect on stiffness perception (Quek et al. 2014). It was also shown that when applying tangential and normal forces to the tip of the finger, the perceived stretch intensity was linearly related to the applied force magnitude (Paré, Carnahan & Smith 2002). In a sensory substitution study, a tactor-induced skin-stretch device was successfully implemented to convey stiffness information in a teleoperated palpation task, and was more accurate than the widely used vibration feedback (Schorr et al. 2015). Skin-stretch has also been used to convey direction (Gleeson, Horschel & Provancher 2009), augment friction (Provancher, Sylvester 2009), and replace kinesthetic information in navigation tasks (Quek et al. 2015a). These findings clearly establish that stretching the skin of the finger pads augments perception, but they do not provide any information about the role of skin-stretch information in the control of grip force.

During tool-mediated interactions with objects, humans apply grip force perpendicularly to the held object to prevent its slippage or breakage. Three known components contribute to grip force when interacting with an object: a predictive feedforward component that consists of a baseline grip force and a modulation of grip force with anticipation of the load force, and a reactive feedback component that responds to slippage. Baseline grip force is maintained to create a safety margin, and is increased in the case of uncertainty about the load force (Gibo, Bastian & Okamura 2014). Feedforward control is used to adjust the grip force in accordance with the expected slipperiness and weight of the object (Flanagan et al. 2003), or any load force (Flanagan, Wing 1990, Flanagan, Wing 1997, Danion, Sarlegna 2007, Danion, Descoins & Bootsma 2009). It is widely accepted that the nervous system can learn and form internal representations of object dynamics (Johansson, Cole 1992, Shadmehr, Mussa-Ivaldi 1994, Kawato 1999, Davidson, Wolpert 2004). Finally, if cutaneous receptors indicate that slippage is occurring, the grip force is increased through rapid feedback control (Kandel et al. 2000, Johansson, Flanagan 2009).

What happens to the control of grip force when tactile or kinesthetic information is missing? During digital anesthesia, participants applied higher grip force, but the anticipatory temporal coupling between the grip force-load force was not impaired (Nowak et al. 2001). This led to the conclusion that cutaneous information from grasping fingers is only necessary for scaling the strength of the grip force, whereas the timing is predictively regulated by kinesthetic information (Witney et al. 2004). Using rTMS to interfere with the normal function of grip force adjustment, White et al. (White et al. 2013) found that the left supplementary motor area is involved in the scaling of grip force during movement preparation. Gibo et al. (Gibo, Bastian & Okamura 2014) found that load force feedback is crucial for grip force-load force modulation during virtual object interactions. In the complete absence of force information, grip force was no longer modulated with load force; rather, the baseline grip force increased to prevent slippage. However, the design of this study made it impossible to separate out the cutaneous and kinesthetic contributions. Thus overall, most of the studies have been able to explore the contribution of the different force sensing modalities to the different components of grip force by eliminating the feedback, disrupting the control, or studying impaired patients.

In contrast, the tactile stimulation devices provide the opportunity to partially disrupt the natural coupling between tactile and kinesthetic information, and dissociate the effects of tactile and kinesthetic channels from the human control of grip force when all the sensory and motor components are intact and not anesthetized. Two recent studies using this technology yielded conflicting evidence on the effect of skin-stretch on grip force: in (Quek et al. 2015a) the mean grip force increased due to skin-stretch, whereas in (Quek et al. 2015b) the mean grip force was not affected. These two studies however only examined the effect of adding artificial skin-stretch on the change in mean grip force, and not on the detailed changes in the different components of grip force as a function of the change in load force and skin-stretch deformation. Hence, the dissociable effect of adding an artificial skin-stretch on the three components of grip force control still remains unclear.

There are many examples in the literature of the dissociation between perception and action during the grasping (Aglioti, DeSouza & Goodale 1995, Haffenden, Schiff & Goodale 2001) and lifting of objects (Flanagan, Beltzner 2000, Bringoux, Lepecq & Danion 2012). The size-weight illusion affects the scaling of grip force in the first trials of a lifting task, but with repeated lifting movements the motor effect fades away, while the illusion persists (Flanagan, Beltzner 2000). In contrast, the introduction of a delay in visual feedback of movement led to a persistent alteration of grip force control that could be accounted by an illusory change in object dynamics (Sarlegna, Baud-Bovy & Danion 2010). Leib et al. (Leib, Karniel & Nisky 2015) found that during interactions with linear elastic force fields, a delay in the force feedback produced the illusion of touching a softer elastic force field, but the participants’ grip force was predictively adjusted to the correct stiffness level and timing. This dissociation could be specific to the delayed feedback, but it could also be related to differences in the contribution of tactile and kinesthetic information. Hence, it remains to be determined whether the perceptual augmentation of stiffness due to tactile stimulation would also evident in the predictive grip force adjustment.

With this question in mind, we designed two experiments to test the effects of adding artificial skin-stretch to kinesthetic force on perception of stiffness and on the predictive and reactive mechanisms of grip force adjustment. While humans can experience various types of kinesthetic forces, here we focused on the load force. This force is commonly experienced during interactions with objects and it depends on the mechanical properties of the object such as its mass, stiffness or viscosity. Specifically, we conducted two experiments in which participants were asked to interact with and judge the stiffness of virtual objects – elastic force fields with different levels of stiffness that were rendered with a kinesthetic haptic device and augmented with different levels of additional stretch stimulation to the skin of the fingertips. In the first experiment, we focused on determining how the additional stretch stimulation of the fingertips biases perception of stiffness and changes the overall trajectories of grip force. In the second experiment, we embedded stretch-catch probed in different stages of the repeated interactions of each elastic force field to dissect the effect of the tactile stimulation on the different components of grip force control.

We hypothesized that due to the additional stimulation of the cutaneous mechanoreceptors by the stretch stimulation, we would observe an increase in the perceived stiffness of the force fields and in the reactive component of grip force control as an outcome of feedback. If perception and predictive control of grip force share mutual stiffness estimation mechanisms, we also expected a greater feedforward grip force that would increase the modulation with anticipation of the load force following repeated interactions with the force field. However, if there is a dissociation between perception and predictive control of grip force, concurrently with increased perceived stiffness, the predictive grip force would not be affected by the stretch. In addition, the skin-stretch stimulus may also increase load force uncertainty, and therefore, participants might increase their baseline grip force to increase the safety margin regardless to the perceptual effect or the scaled modulation.

## Results

### Experiment 1

We used a stiffness discrimination task to examine the contributions of adding artificial skin-stretch to a kinesthetic load force in the formation of stiffness perception and the control of the applied grip force during unconstrained interactions. Participants (N=10) sat in front of a virtual reality setup and interacted with virtual elastic force fields that were rendered with a haptic device [Fig. 1]. In some of the force fields, they also received skin-stretch feedback via a custom-build tactile stimulation device. The magnitude of the stretch was proportional to their penetration depth into the virtual elastic force field, in addition to the load force feedback. In a forced-choice paradigm, we asked participants which of the two force fields had a higher level of stiffness. In each pair, the stiffness of the *comparison* force field was chosen from ten possible values, and the stiffness of the *standard* force field was set to a constant value. During the interaction with the *standard* force field, we stretched the skin of the thumb and index fingers using the skin-stretch device [Fig. 1]. The gain of the skin stretch stimulation was chosen from four possible values (0, 33, 66, and 100 mm/m).

**Fig. 1.**
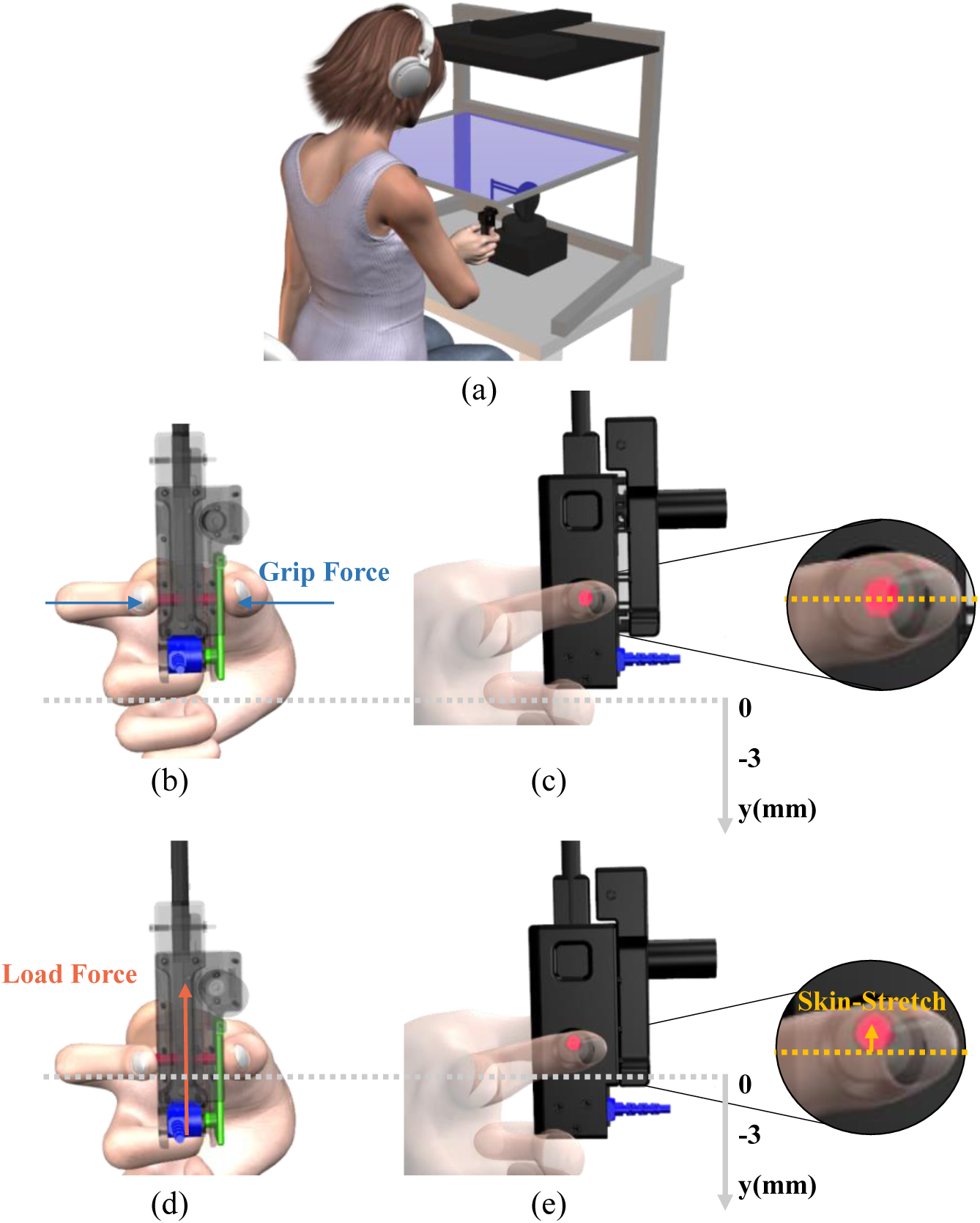
Experimental system. (a) The participants sat in front of a virtual reality rig, and held the skin stretch device that was mounted on a haptic device. (b) Back and (c) side views of the skin stretch device while there was no interaction with the force field, and therefore the load force and the skin-stretch were set to be zero. Zoom-in view of (c) shows that the tactor was in its zero state. (d) Back and (e) side views of the skin-stretch device during interactions with the force field. Both the load force and the skin-stretch increased as the penetration into the force field increased. Zoom-in view of (d) shows that the tactor was moving upwards.

#### Skin-stretch caused perceptual overestimation of stiffness

The addition of artificial skin-stretch to a kinesthetic load force caused participants to overestimate the stiffness of the *standard* force field. Using psychometric functions that were fitted to the probability that the participants would respond that the *comparison* force field was stiffer as a function of stiffness difference between the *comparison* and *standard* force fields, we found that the augmentation in the perceived stiffness increased linearly as a function of the skin-stretch gain. The psychometric curves of a typical participant in Fig. 2(a) show that a higher level of tactor displacement gain caused the illusion of a stiffer force field and thus an overestimation of the perceived stiffness.

**Fig. 2.**
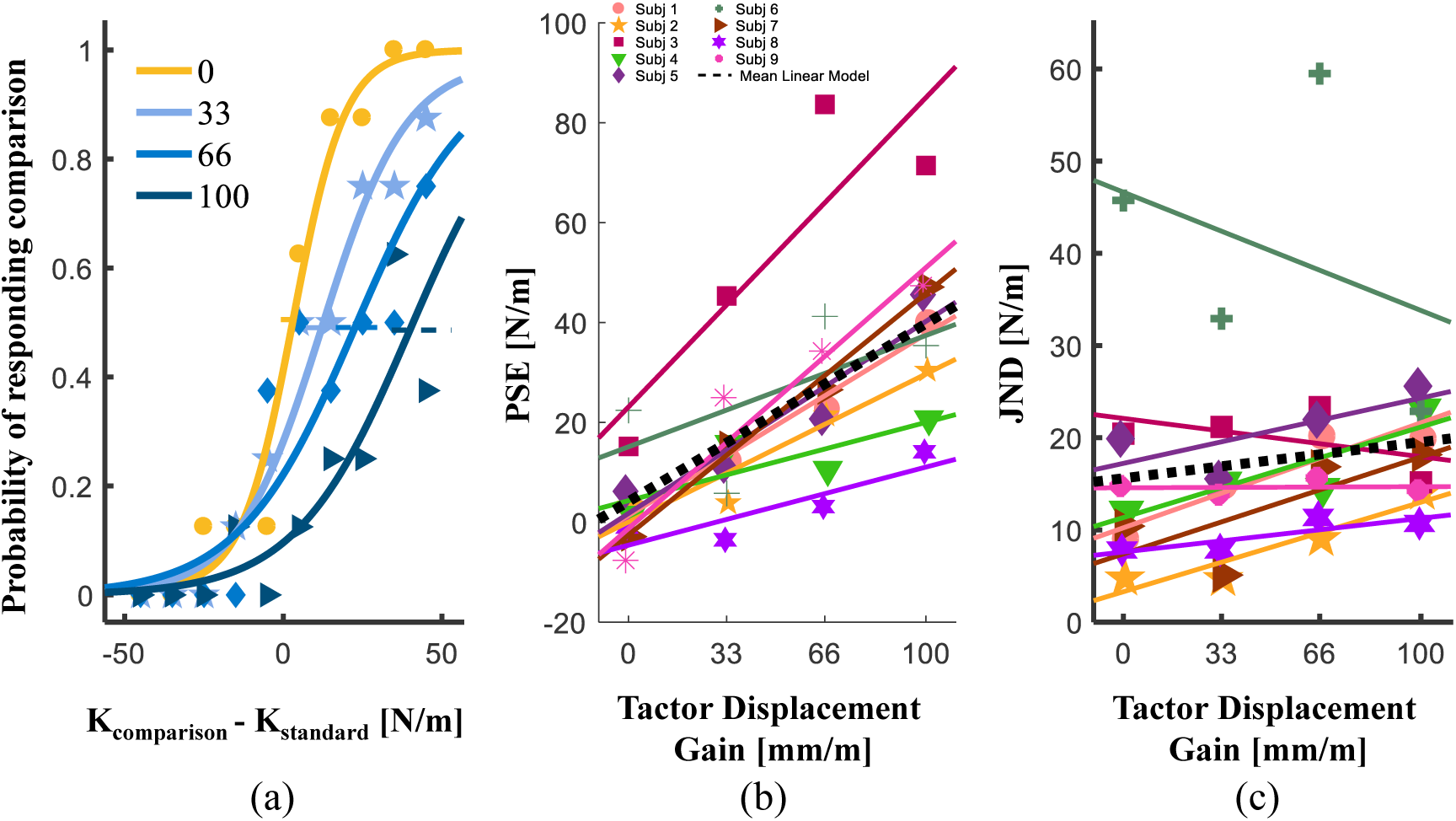
The effect of artificial stretch of the skin on perception. (a) An example of psychometric curves of a typical participant for different levels of skin stretch gain. The abscissa is the difference between the stiffness of the comparison and the standard force fields, and the ordinate is the probability of responding that the comparison force field had a higher level of stiffness. Each line represents the participant’s response for different values of the gain. The horizontal dashed lines represent the standard errors for the PSE. (b) The point of subjective equality values as a function of the tactor displacement gain. (c) The just noticeable difference values as a function of the tactor displacement gain. The markers represent the PSE and JND (b and c, respectively) values for each participant. The solid lines represent the linear regression of each participant, and the black dotted line is the average regression line across all participants.

In trials without artificial skin-stretch of the finger pad; i.e., when the tactor displacement gain was set to zero, the Point of Subjective Equality (PSE, a measure of bias in the perceived stiffness) was close to zero. This means that the participant could accurately distinguish the stiffness of the two force fields. Stretching the finger-pad skin of this participant during the interaction with the *standard force* field caused a rightward shift of the curve and a positive PSE, indicating that this participant overestimated the stiffness of the *standard* force field. Increasing the tactor displacement gain caused an additional rightward shift of the PSE. This shift was observed in nine out of the ten participants [Fig. 2(b)], and was statistically significant (rm-regression, main effect of ‘gain’: *F*_(1,8)_ = 42.47, *p* = 0.0002). The increase in the average perceived stiffness was about 40 N/m; i.e., 47% of the kinesthetic stiffness level. The only participant who showed the opposite trend in perceived stiffness reported during the experiment that she became aware of the applied skin-stretch, and tried to resist its effect by responding in the opposite way to what she felt. Therefore, this participant was excluded from the statistical analysis of perception (which did not affect the statistical significance of any of our conclusions).

In contrast to the bias in stiffness perception (as quantified by the PSE) that was affected by the skin-stretch gain, we did not find a difference in discrimination sensitivity (as quantified by the JND) with an increase in tactor displacement gain (rm-regression, main effect of ‘gain’: *F*_(1,8)_ = 2.060, *p* = 0.189). Some of the participants did show an increase in JND with an increase in tactor displacement gain, but the increase across all participants was not statistically significant [Fig. 2(c)]. Hence, we cannot conclude that the added tactile stimulation impairs the discrimination accuracy.

#### Skin-stretch caused an increase in the applied grip force

Adding the skin-stretch stimulus maintained the grip force-load force modulation [Fig. 3]. Surprisingly, the grip force trajectories had a non-uniform peak pattern that appeared predominantly in trials in which the skin-stretch was applied. For example, the grip force signals in the examples in Fig. 3 show that increasing the skin-stretch gain caused more grip force peaks to appear. In addition, when skin-stretch was applied, there was a rising trend in the magnitude of the applied grip force as the participant made subsequent probing movements (compare the zero gain and the 100 mm/m gain plots). The grip force trajectories in Fig. 3. also create the impression that there are phase differences between grip force responses and the load force due to added tactor displacement. However, another (and more likely in light of our results of Experiment 2) explanation to these changes is a reactive response to the skin-stretch deformation which changes the uniform grip force pattern (see experiment 2 and SI for more information).

**Fig. 3.**
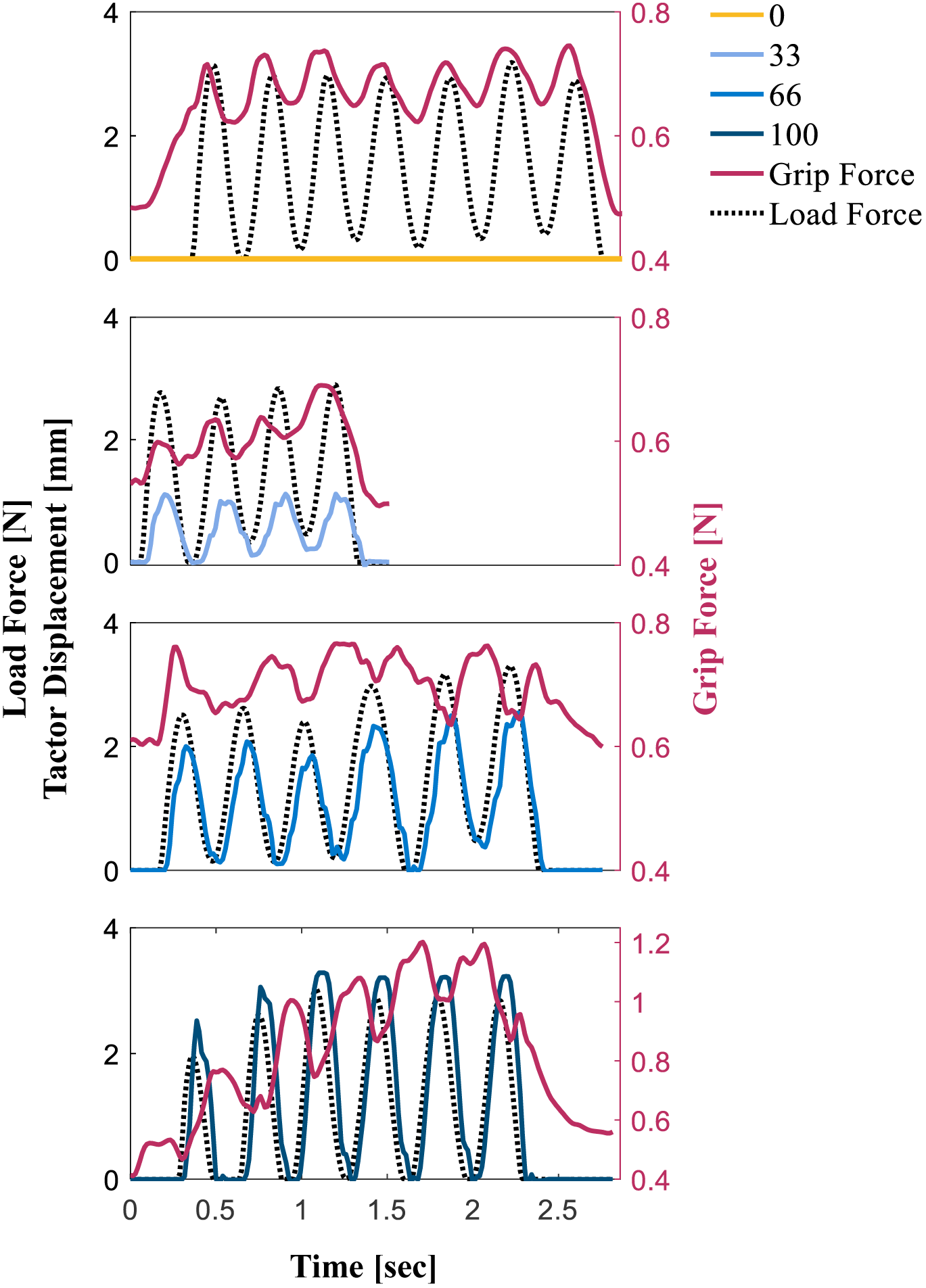
Examples of load force, grip force, and tactor displacement trajectories during typical trials of a typical participant.

To quantify the change in grip force with repeated probing interactions within each trial, we analyzed the *peak grip force-peak load force ratio*, and compared the first to the last probing movements in terms of this ratio. The difference between the last grip force-load force peak ratio and the first grip force-load force peak ratio for each participant is depicted in Fig. 4(a). Consistent with previous studies (Leib, Karniel & Nisky 2015), for the zero tactor displacement gain, this difference was negative; in other words, the participants decreased their grip force as they formed a representation of the force field that they touched. Increasing the tactor displacement gain led to a relative increase in *the peak grip force-peak load force ratio*. This means that participants applied more grip force per amount of kinesthetic load force during the last probing movements when interacting with force fields with higher levels of tactor displacement gain than with lower levels of tactor displacement gain. Fig. 4(b) plots the average ratios across all participants, for each tactor displacement gain on the first and last probing movements. This analysis confirmed that during the last probing movement, *the peak grip force-peak load force ratio* increased as the tactor displacement gain increased. These effects were statistically significant (rm-ANCOVA, interaction between ‘gain’ and ‘probing movement’ variables: *F*_(1,49)_ = 37.65, *p* < 0.0001). The increase in the *peak grip force-peak load force ratio* between the 0 mm/m and 100 mm/m was 0.17, which is 48.5% of the value at 0 mm/m, and comparable to the amount of augmentation in perception.

**Fig. 4.**
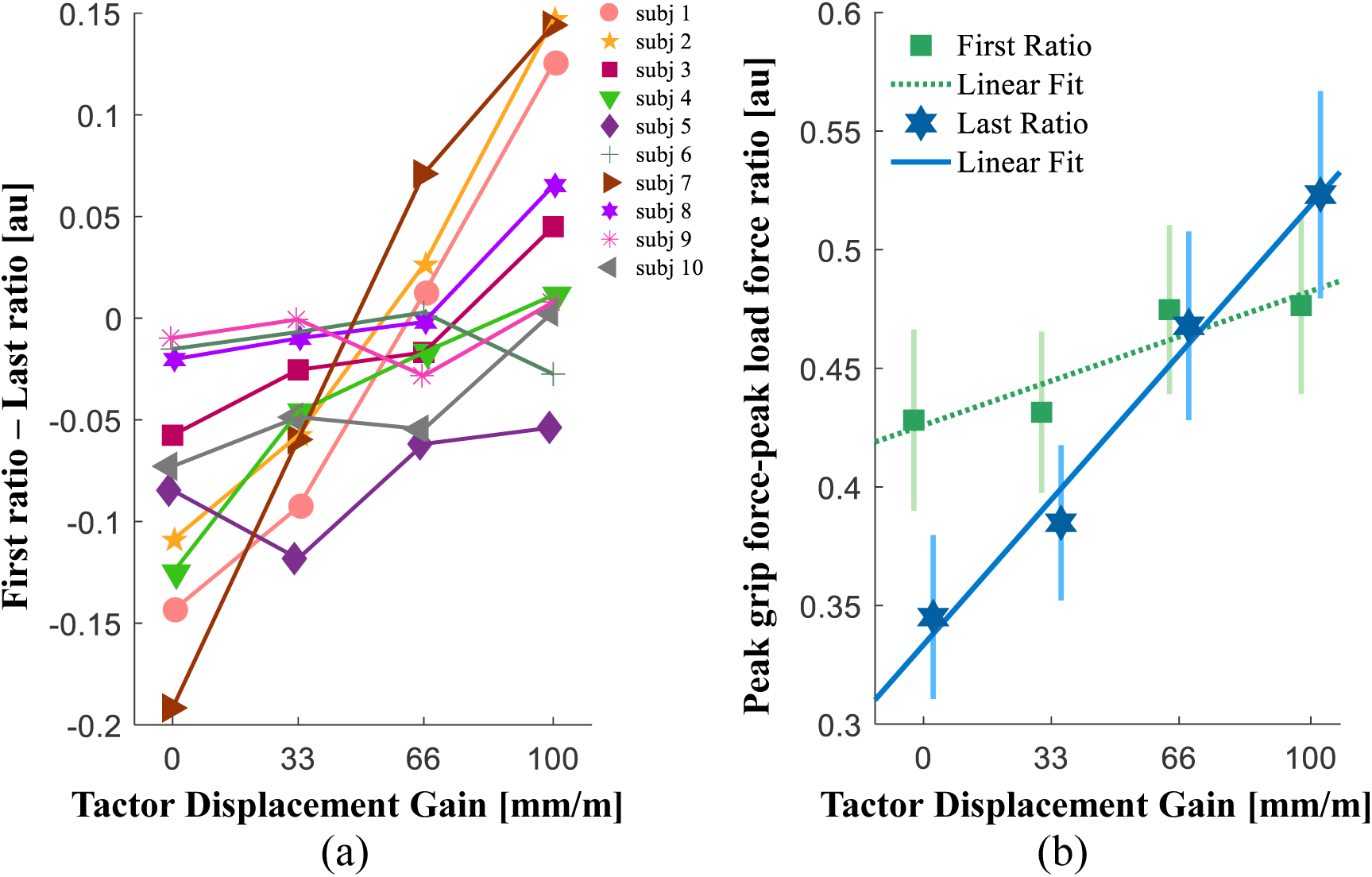
Analysis of peak grip force-peak load force ratio. (a) The difference between the peak grip force-peak load force ratios in the last and the first probing movements for individual participants. (b) The average ratios during the first (green) and last (blue) probing movements. The markers and vertical lines represent the average peak grip force-peak load force ratios across all participants and their standard errors. The dotted green and solid blue lines represent the average fitted regression lines.

The slope of the regression line of the *peak grip force-peak load force ratio* for the first probing movement was much shallower than for the last probing movement, but surprisingly, it was significantly greater than zero (rm-regression, main effect of ‘gain’: *F*_(1,9)_ = 6.63, *p* = 0.0300). Because participants could not predict the tactor displacement gain value at their initial interaction with the force field, it is likely that this increase in grip force was a result of a reactive response. In addition, we also found that in our experimental system, there was a small artifact of tactor displacement on the measurements of the force sensor. This artifact was measured in all tactor displacements without human contact, but its size was much smaller than the size of the effects on grip force that we observed in the entire study (see SI for its characterization).

Overall, in this experiment, we showed that adding artificial skin-stretch increased the perceived stiffness linearly with the tactor displacement gain. We also found that the artificial skin-stretch led to an increase in grip force that developed with subsequent probing interactions with the same elastic force field. However, we could not differentiate the contributions of the different components of grip force; namely, the predictive feedforward component that modulates grip force in anticipation of the load force, the predictive baseline grip force, and the reactive feedback component. Experiment 2 was designed to resolve this issue.

### Experiment 2

This experiment was designed to quantify the dissociable contribution of skin stretch to the three components of applied grip force. We designed a new experiment, similar to Experiment 1, but with stretch-catch probes where we maintained the load force but unanticipatedly omitted the skin-stretch. The participants (N=10) were asked to make exactly eight discrete movements in each of the two force fields. We added stretch-catch probes during the interaction with the *standard* force field in the second and seventh probing movements. This protocol allowed us to differentiate the feedforward and feedback grip force components from the applied grip force. Calculating the difference between the seventh and the second stretch-catch probe of similar trials was used to identify the predictive component of grip force control that accumulates with repeated stretch stimulation. Calculating the difference between similar regular and stretch-catch probes (i.e. with and without artificial skin-stretch) in the second probing movements was used to isolate the reactive component of grip force control.

Fig. 5 presents examples of grip force, load force, and tactor displacement trajectories during the regular and stretch-catch probes. During regular probes with high levels of artificial skin-stretch (gain 66 and 100 mm/m), the grip force trajectories had irregular pattern of decrease accompanied by an increase. Comparing these trajectories with the trajectories of the stretch-catch probes demonstrate the reactive response of grip force to the skin-stretch stimulation with our device. During stretch-catch probes and probes without high levels of artificial skin-stretch (gain 0 and 33 mm/m and during stretch-catch probes), each grip force peak was coupled with a load force peak.

**Fig. 5.**
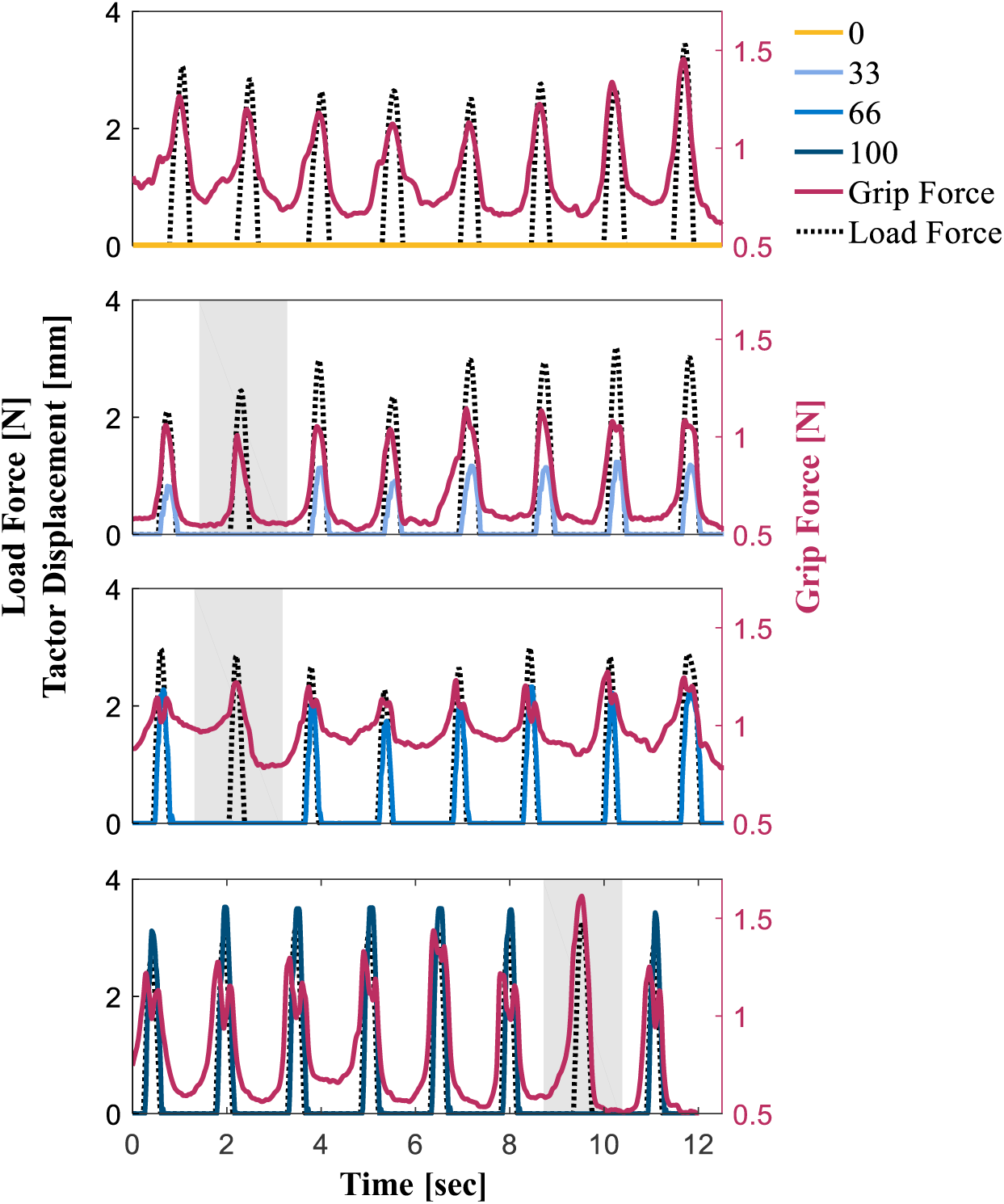
Examples of load force, grip force, and tactor displacement trajectories during typical trials of a typical participant. The gray shaded regions highlight stretch-catch probes.

#### Skin-stretch increased the feedforward component of grip force

Fig. 6 shows the results of the grip force-load force ratio analyses. The grip force trajectories in Fig. 6 were normalized by the peak load force, and were time-normalized and aligned such that 0 was the onset of the contact with the force field, and 1 was the end of the interaction.

**Fig. 6.**
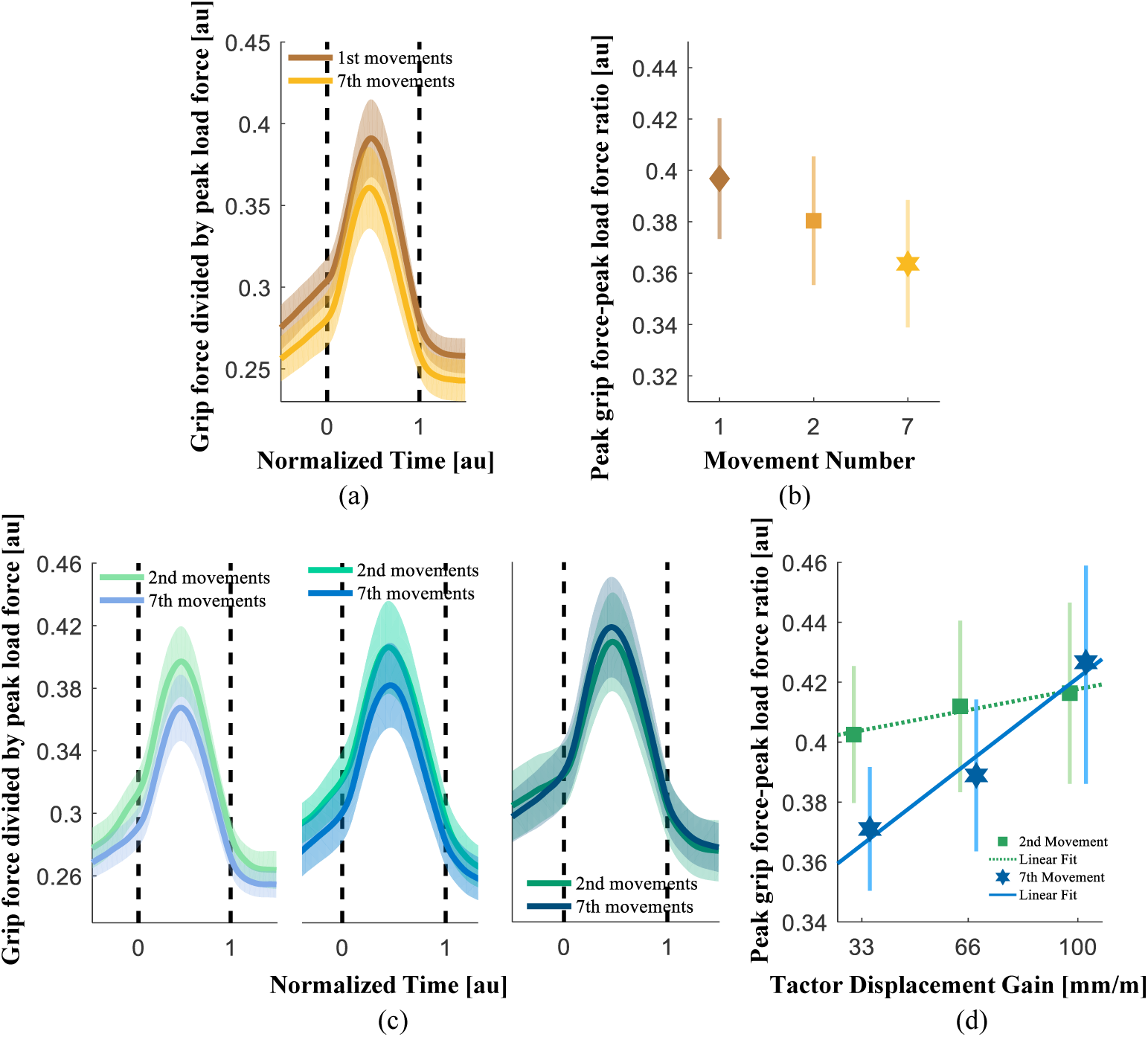
Grip Force-Load Force Ratio. (a) The average grip force trajectories divided by the peak load force across all participants, for the first (brown) and seventh (orange) probing movements. (b) Analysis of peak grip force-peak load force ratio during the first, second, and seventh probing movements. (c) The average grip force trajectories divided by peak load force across all participants, for the second (green) and seventh (blue) probing movements (left: 33 mm/m, middle: 66 mm/m, and right: 100 mm/m). (d) Analysis of the peak grip force-peak load force ratio during the second (green) and seventh (blue) probing movements. In (a) and (c) the shading represents the standard errors, and the black vertical dashed lines represent the onset and the end of the contact with the elastic force field. In (b) and (d) the markers and the vertical lines represent the average peak grip force-peak load force ratios across all participants, and their standard errors. The dotted green and solid blue lines represent the average fitted regression lines.

In trials without skin-stretch [Fig. 6(a) and (b)]; i.e., with a gain of 0 mm/m, the *peak grip force-peak load force ratio* decreased between the first and seventh probing movements (rm-regression, main effect of ‘probing movement’: *F*_(1,9)_ = 10.97, *p* = 0.0090), and also between the second and seventh probing movements (rm-regression, main effect of ‘probing movement’: *F*_(1,9)_ = 6.69, *p* = 0.0294).

Unlike Experiment 1, in trials with a positive tactor-displacement gain, to examine the effect of skin stretch only on the predictive component of grip force, we only analyzed the grip force-load force trajectories of the stretch-catch probes. Fig. 6(c) presents the average grip force trajectories divided by the peak load force as a function of tactor displacement gain, on the second and seventh probing movements. We observed that the average grip force profiles decreased between the second and seventh probing movements, but increased as the gain value increased. The increase with gain value was evident for both the second and seventh probing movements, but this increase was greater for the seventh probing movement. Fig. 6(d) shows the rising trend in the applied grip force with higher levels of tactor displacement gain for both the second (gentle rise) and seventh (steep rise) probing movements (rm-ANCOVA, interaction between ‘gain’ and ‘probing movement’ variables: *F*_(129)_ = 8.88, *p* = 0.0058).

These results are similar to those described in Experiment 1, but here were only measured in the predictive grip force component. The gentle increase in the second probing movement can be explained by the fact that participants had already been exposed to the skin-stretch stimulus in the first probing movement, and could generate a prediction of the stimulus.

This analysis however, does not dissociate between the two components of the predictive grip force control - the baseline that does not incorporate a specific representation of the force field, and the modulation in anticipation of the load force. To separate these two components, we ran a linear regression analysis in the grip force-load force plane. The intercept of this regression represents the baseline grip force; i.e., - the amount of grip force when no external load force is applied by the haptic device. The slope of the regression represents the grip force-load force modulation. In typical manipulation tasks of grasping and lifting, the slope is usually linked to the “slip ratio” of an object, but in our case, the increase in the slope represents the motor illusion of a harder object due to skin stretch.

In trials without skin-stretch; i.e., with a gain of 0 mm/m, the intercept and the slope decreased between the first and second probing movements (rm-regression, main effect of ‘probing movement’: slope - *F*_(1,9)_ = 6.15, *p* = 0.0350, intercept - *F*_(1,9)_ = 8.81, *p* = 0.0158), but did not change between the second and seventh probing movement [Fig. 7(a) and (b)]. In contrast to the previous analysis, here we did not add 50 samples before and after the start and end movement points, but only examined the forces that were applied during the interaction with the force fields. This change can help account for the difference in the results for the two different analyses of zero stretch [Fig. 6(b) and Fig. 7(a) and 7(b)]. Fig. 7(c) shows that the intercept increased with an increase in the tactor displacement gain (rm-ANCOVA, main effect of ‘gain’: *F*_(1,9)_ = 8.03, *p* = 0.0196), but surprisingly, we did not find a difference between the second and seventh probing movements (rm-ANCOVA, interaction between ‘gain’ and ‘probing movement’: *F*_(1,29)_ = 0.50, *p* = 0.4848). The analysis of gain zero hints that there may have been a rapid change in the applied grip force between the first and second probing movement which would explain why we did not find a difference between the second and seventh probing movements.

**Fig. 7.**
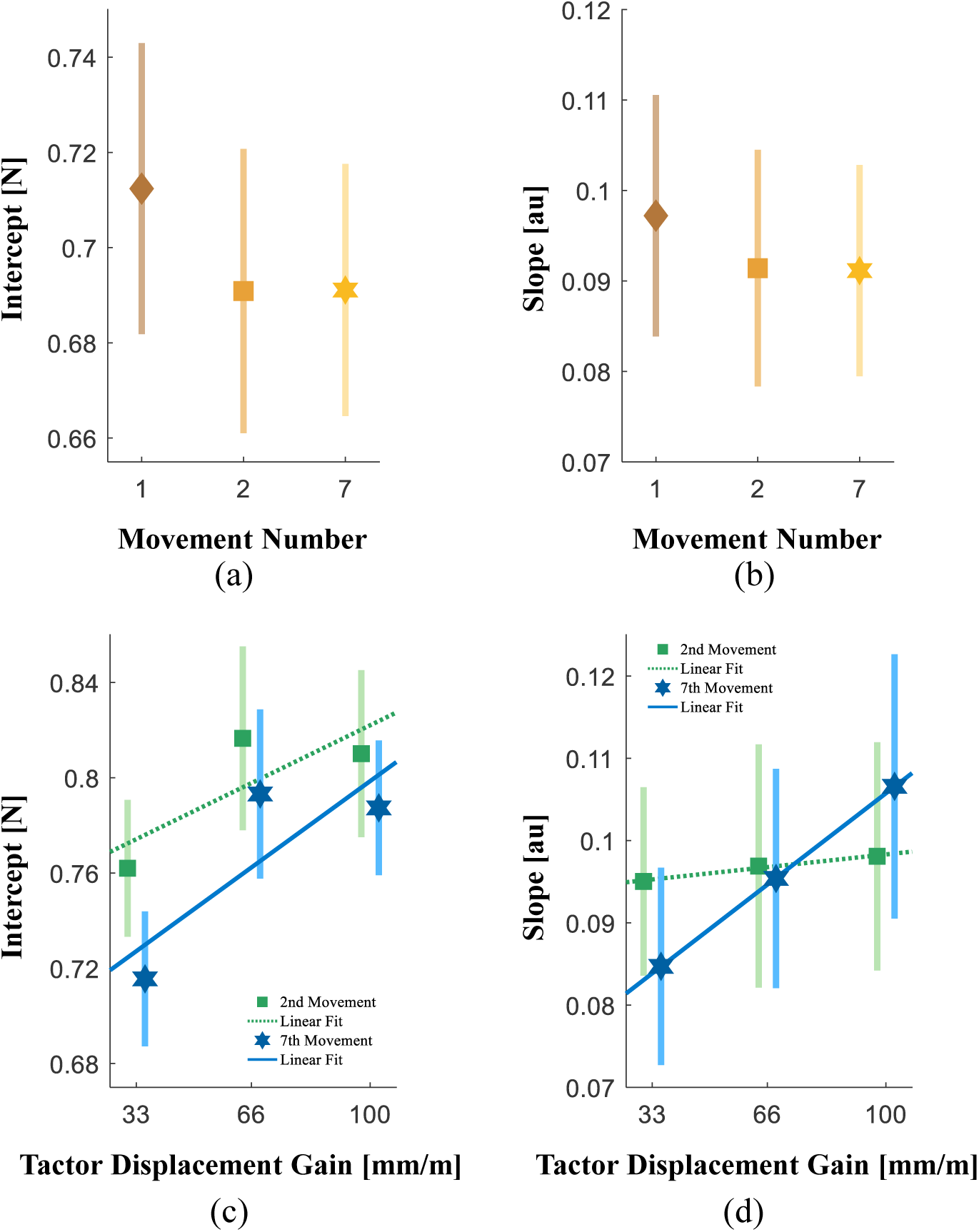
The regression analysis parameters, (a) Intercept and (b) slope for a gain of 0 mm/m, in the first, second, and seventh probing movements. (c) Slope and (d) intercept, for different levels of tactor displacement gain. The dotted green and solid blue lines represent the average fitted regression lines. The markers and the vertical lines represent the average values across all participants and their standard errors.

Fig. 7(d) shows that the slope of the second probing movement did not change across different tactor displacement gains, but the slope of the seventh probing movement increased with an increase in tactor displacement gain (rm-ANCOVA, interaction between ‘gain’ and ‘probing movement’: *F*_(1,29)_ = 5.13, *p* = 0.0311). This analysis suggests that although the intercept did not change during repeated interactions with the force field, the slope changed as function of tactor displacement gain. It is important to mention that the slopes values are small with respect to previous studies (Flanagan, Wing 1990, Leib, Karniel & Nisky 2015) because we measured a downscaled version of the grip force (see the methods section for more information).

*The peak grip force-peak load force ratio* analysis of skin-stretch gains (33, 66, and 100 mm/m) revealed a decrease between initial and late exposure (rm-ANCOVA, main effect of ‘probing movement’: *F*_(1,34)_ = 10.10, *p* = 0.0032). However, we did not find a change in either the intercept or the slope between the initial and late exposure in the *grip force load force regression analysis* (main effect of ‘probing movement’: slope - *F*_(1,35)_ = 3.71, *p* = 0.0623, intercept - *F*_(1,35)_ = 2.38, *p* = 0.1316). We believe that the significant decrease that was observed in the first analysis [Fig, 6(d)] can be attributed to an additive effect of the moderate decreases of the two components observed in the second analysis [Fig. 7(c) and (d)].

#### The effect of skin-stretch on the feedback component of grip force control

To isolate the reactive component of grip force control, we calculated the difference between regular and stretch-catch probes (i.e. with and without artificial skin-stretch) in the second probing movement. Because of the close proximity between the tactor displacement and the measurement of grip force, there was an artifact of the tactor movement in our grip force measurements (see SI for its identification).

Fig. 8(a) and (b) present the reactive response before and after subtracting the artifact of tactor displacement on the measured grip force, respectively. The grip force trajectories in Fig. 8 were normalized by the peak load force, and were time-normalized and aligned such that 0 was the onset of the contact with the force field, and 1 was the end of the interaction. Surprisingly, visual examination suggests that the stretch of the skin caused a reactive decrease in grip force; this decrease was larger for larger tactor displacement gain. In addition, at the end of the interaction, participants increased their grip, and the magnitude of the increase was also larger for larger tactor displacement gains. Overall, the difference between the reactive grip force at the end of the interaction with the elastic force field and the minimum of the grip force trajectory (*the reactive grip force range*) depended linearly on the tactor displacement gain (rm-regression, main effect of ‘gain’: *F*_(1,9)_ = 28.37, *p* = 0.0005). This pattern of the reactive response is likely responsible for the irregular peak pattern that we observed in Experiment 1 and in Fig. 3. It is important to acknowledge that the reactive grip force response we report here may be a specific response to our skin-stretch device.

**Fig. 8.**
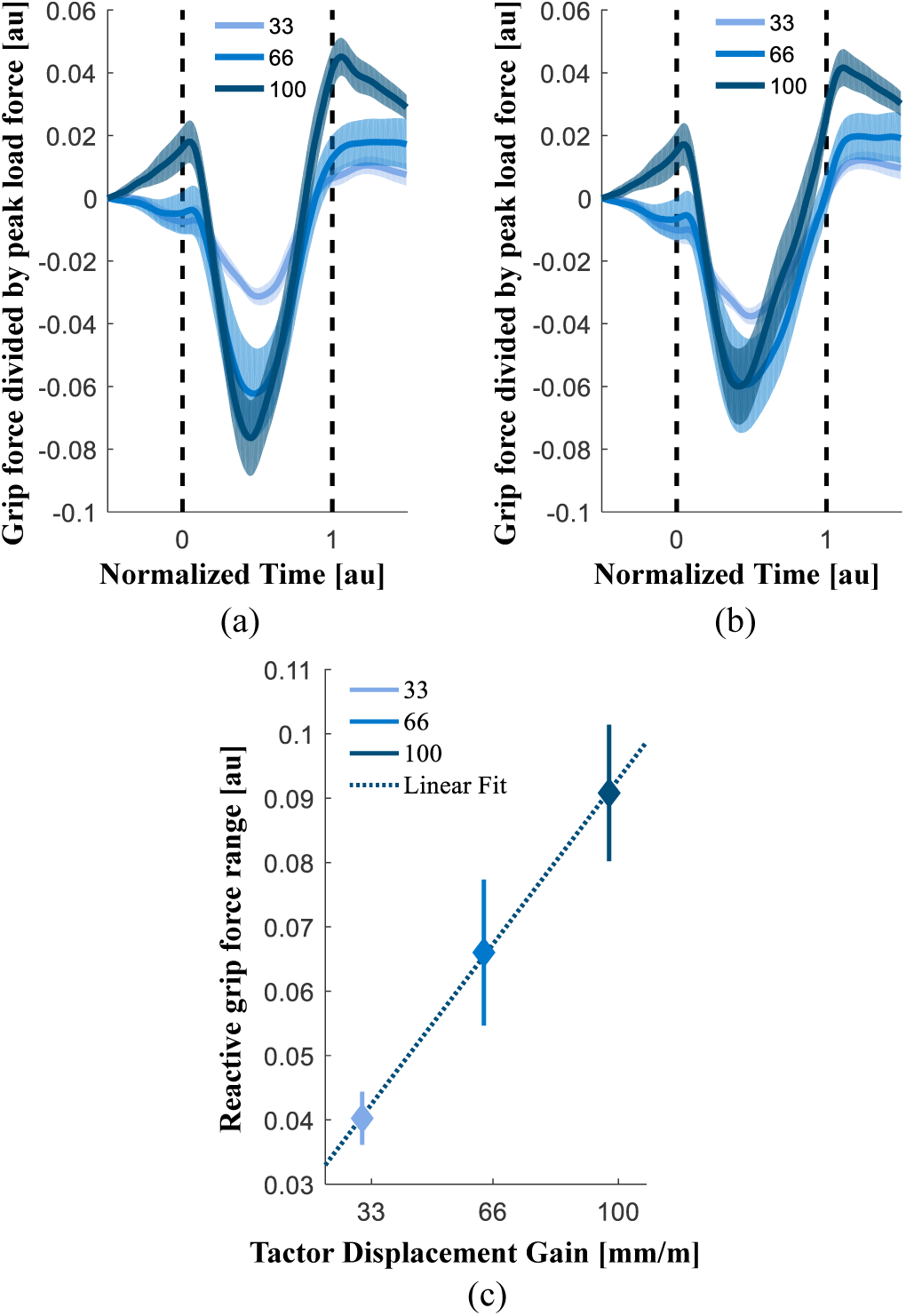
Reactive grip force component. This component was calculated by subtracting the second stretch-catch probing from the second probing movement with both skin-stretch and load force stimuli. The trajectories depict grip force trajectories divided by the peak load force. (a) Participants’ reactive responses together with the artifact. (b) The reactive response after subtraction of the artifact. Shading represents the standard errors, and the black vertical dashed lines represent the time when the participants interacted with the elastic force field. (c) The reactive grip force range; the difference between the reactive grip force at the end of the interaction with the elastic force field and the minimum of the grip force trajectory. The blue squares represent the average values across all participants, and the vertical lines represent the standard errors.

Thus, in this experiment, we found that high levels of artificial skin-stretch caused an increase in grip force-load force modulation that developed with subsequent probing interactions with the same elastic force field. Because of the artificial skin-stretch, participants did not reduce the safety margin even after an internal representation was formed. In addition, we showed that *the reactive grip force range* also depended linearly on the tactor displacement gain.

## Discussion

In this study, we examined how kinesthetic force in the form of a load force, and artificial tactile feedback in the form of a skin-stretch, affect the perception of stiffness and cause grip force adjustments during interactions with viscoelastic objects. Our results suggest that adding an artificial skin-stretch to a kinesthetic force creates the illusion of interacting with a stiffer object and this illusion also affects how participants act upon the object. In both action and perception, the change was linearly related to the intensity of the applied stretch, as expressed by the displacement gain of the tactor, and the relative size of the effect was similar. In the first experiment, we focused on the tactor displacement-induced perceptual and motor illusion in unconstrained interactions with elastic force fields. In the second experiment, we focused on dissecting the effect of the tactor-induced illusion on the different components of grip force control. We found that the increase in grip force is reflected in both predictive and reactive components. The predictive grip force was modulated to an extent consistent with a tactile-induced illusion of an increased load force. Unlike previous studies that have examined the dissociable contribution of cutaneous and kinesthetic information through anesthesia or impairment, we addressed this question when both channels were intact.

Consistent with previous studies (Quek et al. 2014, Quek et al. 2013), we found that adding artificial tactile feedback in the direction of the applied kinesthetic force augmented the perceived stiffness, and that the augmentation was a linear function of tactor displacement gain. Nine out of the ten participants showed an increase in perceived stiffness, but there was variability in the magnitude of the effects. This variability could be due to variations in participants’ mechanical skin properties (Wang, Hayward 2007), the way that participants held the device, and/or the amount of grip force they applied. Participants who held the skin-stretch device with a higher level of initial grip force could feel the skin-stretch stimulus more strongly and therefore their perceptual illusion was greater. This could be investigated in future studies by measuring the displacement of the skin finger pad using the method described in (Delhaye, Lefevre & Thonnard 2014) but is outside of the scope of our current study.

Our results resemble those reported in Quek et al. (Quek et al. 2014), who showed that inducing artificial skin-stretch leads to an increase in the perceived stiffness of kinesthetically rendered surfaces. However, in their study, participants also received visual feedback, which may explain the large inter-subject variability that was less prominent in our results. By eliminating the visual feedback, we highlighted the contribution of haptic feedback in perceptual assessment of the stiffness. The augmentation effect we found is consistent with previous studies which reported that adding tactile feedback to kinesthetic forces augments perception of friction (Provancher, Sylvester 2009) and force (Matsui, Okamoto & Yamada 2014).

Different studies have aimed to find the computational basis for cognitive estimations of stiffness (Chib et al. 2009, Nisky et al. 2014, Casadio, Pressman & Mussa-Ivaldi 2015). One such example is a model that accounts for the formation of stiffness estimations as a function of a combination of estimations of environment stiffness (impedance) and the inverse of environment compliance (admittance) (Nisky, Mussa-Ivaldi & Karniel 2008, Nisky, Baraduc & Karniel 2010). This means that the sensorimotor system uses the information gathered during probing movements to form a stiffness judgment as a combination of two estimations: the ratio between sensed force and the desired/resultant hand position (impedance) and the ratio between sensed hand position and the desired/resultant force (admittance). Each estimation is weighted in the general stiffness judgment depending on the way the probing movement is performed, such as the fraction of probing movements where the participant broke contact with the force field (Nisky, Mussa-Ivaldi & Karniel 2008), or the dominant joint used during interaction with the force field (Nisky, Baraduc & Karniel 2010). In this sense, a larger perceived stiffness suggests that the skin acts as a biological strain gauge, and therefore transmits indications of a greater force during increased displacement of the skin, this causes the value of the impedance to increase and the value of the admittance to decrease. Alternatively, if the force would be considered as the controlled variable and the skin stretch as indicator of displacement, the perceived stiffness would be smaller. Our results that participants overestimated the stiffness of the force field during an artificially increased stretch of the skin support the idea that participants estimated the stiffness of the force field as an impedance.

Unlike many studies which have reported that the motor system is robust to perceptual illusions (Aglioti, DeSouza & Goodale 1995, Flanagan, Beltzner 2000, Leib, Karniel & Nisky 2015, Goodale, Milner 1992, Ganel, Goodale 2003), we did not find a dissociation between perception and action. This is consistent with other studies that reported similar findings of lack of dissociation (Franz et al. 2000, Jackson, Shaw 2000, Bruno 2001, Buckingham, Goodale 2010). The average effects across participants in our study are in the same direction of interaction with a harder spring: both the perceived stiffness and grip force control increased with increasing tactor displacement gain. Even though we did not quantify weighting explicitly (Ernst, Banks 2002), based on the sizes of the average effects on perception and action, the tactile and kinesthetic information appear to have been weighted similarly when updating the internal representation used for the perception of stiffness and the grip force adjustment. If indeed these processes are related, this would suggest that there is a common stiffness estimation mechanism for perception and grip force control which is distinct from the control of probing movements (Leib et al. 2016). This view is consistent with evidence that the adaptation of grip forces to perturbing forces most likely involves different mechanisms from adaptation to manipulation forces (Danion, Diamond & Flanagan 2013). However, our experiment could not determine whether perception and grip force control indeed share the same stiffness estimation mechanism or have separate mechanisms which caused simultaneous increase in stiffness perception and applied grip force. For example, while stiffness perception mechanism is likely to be cognitively mediated, the predictive grip force control mechanism is likely an automatic process. Future studies are needed to establish whether and how these processes are linked.

In this study, we dissociated the contribution of tactile stimulation on the predictive component of grip force adjustment in an intact system. Many studies have investigated patients with impairments (Witney et al. 2004, Nowak, Hermsdörfer 2003, Nowak et al. 2003) or digital anesthesia (Nowak et al. 2001, Augurelle et al. 2003) to better understand which information is used to build an internal representation that allows for predictive grip force control. These studies tend to conclude that impaired cutaneous feedback increases overall grip force, but does not break the modulation of grip force in anticipation of load force. Our results complete this picture by showing that artificially applied skin-stretch increased the predictive grip force by increasing both the baseline grip force and the modulation with predicted load force. Unlike these previous studies, the artificial skin-stretch also made it possible to determine that the increase was linear with the amount of stretch.

The predictive grip force-load force modulation as quantified by the slope coefficient changed as function of tactor displacement gain. This finding suggests that participant changed their internal representation, but our study design could not differentiate what changed in the representation. The illusion could be related to an additional load force (Quek et al. 2014, Provancher, Sylvester 2009), or alternatively, the internal representation of the force field’s stiffness could have been updated directly. Johansson et al. (Johansson, Westling 1984) showed that during a lifting task, the slope in the grip force-load force plane increased with surface slipperiness. Flanagan et al. (Flanagan, Wing 1990) reported that both the overall grip force and the extent of grip force to load force modulation increased in contact with smooth surfaces compared to rough surfaces. Therefore, another possible explanation for the increased grip force is that the motor system of the participants could have interpreted the additional tactile stimuli as indication of a more slippery contact surface, which would have thus been entirely unrelated to the perceptual illusion of a harder spring.

Previous studies have shown that to reduce the risk of slippage, participants initially apply excessive grip force and combine a safety margin beyond a sufficient amount of grip force. However, with repeated interactions with the environment, the baseline grip force lessens and adjusts to the expected load force (Leib, Karniel & Nisky 2015). Surprisingly, we did not observe this type of decrease in the baseline grip force (which is represented by the intercept coefficient), even though there was an increase in the modulation of grip force. This means that participants did not reduce the safety margin even after an internal representation was formed. They increased the baseline grip force with higher levels of tactor displacement gain but did not change it from the initial to the late exposure.

There are two possible explanations for this result. The intercept analysis of zero gain suggests that there was a rapid response involving a reduction in the baseline grip force between the first and the second probing movement. This is consistent with the faster adaptation of grip forces compared to manipulation forces (Flanagan et al. 2003). Therefore, in trials with tactile-induced illusion, participants could also have changed the average grip force between the first and the second probing movement such that there was no statistically significant difference between the second and the seventh probing movement. The second possible explanation is that the stretch stimulation may have increased uncertainty about the cutaneous information. This explanation is consistent with the increased JND that we observed for some of the participants indicating worse discrimination sensitivity (as quantified by the JND) with higher levels of tactor displacement gain. A recent study found that grip forces are adapted to average load forces and their variability, whereas manipulation forces are only adapted to average forces (Hadjiosif, Smith 2015). Moreover, uncertainty favors increase in impedance (Franklin et al. 2003, Tee et al. 2010) that may also be associated with a rapid increase in grip force (Tsuji et al. 1995). Therefore, increased uncertainty could have prevented the participants from reducing their baseline grip force.

The role of mechanoreceptors in motor function has been thoroughly investigated over the years. Srinivasan et al. (Srinivasan, Whitehouse & LaMotte 1990) showed that grip force control is important to assure stable holds of objects, and it was suggested to be a result of feedback about slip from cutaneous mechanoreceptors. Johansson et al. (Johansson, Westling 1987) provided evidence that tactile afferents play an important role in adapting the ratio between grip force and load force to the friction between the object and the skin. They suggested that the responses of rapidly adapting type 1 (RA1) mechanoreceptors adjust the grip force to the friction. RA1 mechanoreceptors are critical for grip force control; a slippery surface decreases the excitation threshold thus causing these mechanoreceptors to fire more rapidly, which results in a firmer grip that prevents the object from slipping during unexpected perturbations (Johansson, Flanagan 2009). Our study did not implement neurophysiological methods, but future studies using the skin stretch device in combination with these methods could shed light on the contribution of these different mechanoreceptors to grip force control.

The reactive grip force pattern was very surprising. In the beginning of an interaction, before an internal representation is formed, a tactile stimulus is similar to any mechanical perturbation. It is well-established in the literature that during unexpected load force perturbations, the grip force is automatically adjusted (Cole, Abbs 1988, Johansson, Häger & Bäckström 1992, Cole, Johansson 1993). Recording from tactile afferents during trapezoidal load force perturbations to the digit with different rates of loading (Macefield, Häger-Ross & Johansson 1996) revealed that the mean firing rate was scaled by the slope of the loading ramp. This prompted our initial prediction that the reactive response would be an increase in grip force 50-90 msec after the stimulation (Crevecoeur et al. 2016, Crevecoeur et al. 2017). Instead, in response to the artificial stretch, participants first strongly decreased and only then increased their grip force, resulting in overall moderate increase that depended linearly on the tactor displacement gain. The peak of the decrease was roughly in the middle of the probe, close to the peak of the stretch and the load force, and about 150 msec from the onset of the stretch. The peak of the increase was after the stretch and load force returned to zero, about 300 msec from the onset of the stretch. The initial release might have been the result of an unpleasant or painful interaction, but none of our participants indicated any discomfort. It is important to note that these results should be viewed with caution because of the resemblance between the reactive grip force trajectory and the artifact of tactor movement. However, the participants’ reactive response was much greater than the artifact which strongly implies that the artifact is not a plausible explanation for this response. Future studies, probably with other designs of tactile stimulation devices, are needed to pinpoint the origin of this surprising reactive pattern.

Our findings are applicable to the development of intelligent controllers for robotic hands, wearable finger-grounded tactile haptic devices, and when developing novel technologies for providing haptic information in human-robot physical interactions (Prattichizzo et al. 2013, Pacchierotti, Tirmizi & Prattichizzo 2014, Schorr, Okamura 2017). In most medical robotics applications including robot-assisted surgery (Meli, Pacchierotti & Prattichizzo 2014), assistive devices or prosthetics (Akhtar et al. 2014), the user is deprived of haptic information. In other cases, the haptic information is distorted or has a low gain and quality. Understanding the processing of kinesthetic and tactile information and its integration can lead to improvements in these human-robot interaction applications by providing users with a sense of touch that is tailored to their natural information processing strategies.

Understanding the dissociable effects of kinesthetic and tactile cues on grip force control and the integration between these modalities are critical for the enhancement of feedforward and feedback models, and multisensory integration. In this study, we clearly separated the contribution of stretch of the skin to the predictive and reactive components of grip force control. The findings suggest that stretch of the skin contributed to an increase in the amplitude of the predictive adjustment, but at the same time caused a reactive decrease in grip force that was followed by a reactive increase. These results are important for understanding the remarkable human ability to gracefully manipulate a variety of objects manually without breaking or dropping them, and for developing new technologies simulating the human sense of touch.

## Methods

### Experimental Setup

#### Skin Stretch Device

The goal of this study was to better understand the contribution of adding artificial skin-stretch to a kinesthetic force feedback in the formation of stiffness perception and grip force modulation. To do so, we disentangled the natural relationship between the kinesthetic and tactile information sources using a custom-built 1 DoF skin-stretch device [Fig. 1] which allowed us to stretch the user’s skin to different magnitudes. Placing the thumb and the index fingers in the designated locations allowed us to stretch the finger pads through tactor displacement. Our skin-stretch device was based on the design in (Quek et al. 2014) with several modifications. Our device was equipped with a DC micro motor (Faulhaber, series 1516-SR), a spur gearhead (Faulhaber, series 15/8 with gear ratio of 76:1), an encoder (Faulhaber, series IE2-1024), and an analog motion controller (Faulhaber, series MCDC 3002). We integrated a force sensor (ATI, Nano17) to measure the applied grip force. The force sensor was mounted on the lower end of the device such that participants did not place their fingers directly above the force sensor. The left side of the outer shell was comprised of a ‘door’ with an axis on its upper end, and a cylindrical protrusion facing the force sensor. When the device was held with the index finger and thumb on the apertures, the protrusion pressed the force sensor and the relative grip force was measured. This design did not allow us to measure the exact grip force, but through the law of conservation of angular momentum, we measured a downscaled version of the grip force. The force sensor enables the measurement of change in grip force, and how it is effected by tactor displacement gain. The weight of the skin-stretch device was approximately 200g and it compensated by a weigh that was mounted on the haptic device.

#### System Setup

The participants were seated in front of the virtual reality system and held the skin-stretch device that was mounted on a PHANTOM^®^ Premium 1.5 haptic device (Geomagic) with the index finger and thumb of their dominant right hand [Fig. 1]. The participants looked at a semi-silvered mirror showing the projection of an LCD screen placed horizontally above it. An opaque screen was fixed under the mirror to block the view of the hand. During the experiment, the participants wore noise cancelling headphones (Bose QuietComfort 35 QC35) to eliminate auditory cues from the motor.

We used the haptic device to apply a kinesthetic virtual elastic force field in the vertical upward direction [y in Fig. 1]. The kinesthetic force and the skin-stretch were applied along the same direction, and only after participants were in contact with the force field. Participants could make and break contact with the force field by moving their hand along the vertical axis, and they did not experience any force or stretch while moving along the positive half of the y-axis. While moving along the negative half of the axis, participants experienced force and stretch that were proportional to the amount of penetration distance.

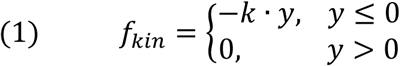

where k [N/m] is the stiffness, and y [m] is the penetration distance into the virtual force field. We used the skin-stretch device to apply tactile stimuli by means of tactor displacement:

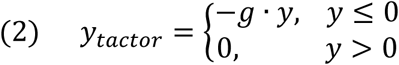

where g [mm/m] is the tactor displacement gain, and y [m] is the same penetration distance into the force field.

#### Protocol

The participants were asked to probe pairs of virtual elastic *standard* and *comparison* force fields and indicate which force field was stiffer. Each force field was indicated to the participants as a screen color that was either red or blue, which we defined randomly prior to the experiment. After the participants interacted with the force fields and decided which one was stiffer, they pressed a keyboard key that corresponded to the screen color of the stiffer force field. For each trial, the stiffness of the *comparison* force field was also pseudo-randomly chosen prior to the experiment. During the interaction with the *comparison* force field, only kinesthetic force was applied. The interaction with the *standard* force field was augmented with one of four different levels of skin-stretch gain (0, 33, 66, and 100 mm/m), in addition to the applied load force. After choosing the force field that they perceived as stiffer, the screen became grey, and to initiate the next trial participants were instructed to stand on a green square. There was no visual feedback along the vertical axis during the experiment. To avoid force saturation of the robotic device, we used an auditory alert if the penetration into the force field exceeded 40 mm. To avoid saturation of the skin-stretch device, we limited the range of the tactor-displacement gains that we investigated to 100 mm/m. This assured that during typical interactions with the force fields, the tactor did not reach the aperture of the device body, and caused as few saturation cases as possible.

### Experiment 1

Ten participants (six females, 22-26 years old) participated in the experiment, after signing an informed consent form as approved by the Human Subject Research Committee of Ben Gurion University of the Negev, Be’er-Sheva, Israel.

In this experiment, the *standard* force field had a constant stiffness value of 85 N/m. During interaction with this force field, the tactor displacement gain was either 0, 33, 66, and 100 mm/m. We chose the *comparison* force field stiffness from a range of values between 40-130 N/m spaced at intervals of 10 N/m. Participants could probe the force fields as many times as they desired and switch between the force fields to make their decision. To switch between the two force fields, participants had to exceed at least 30 mm beyond the boundary of the force field by raising the robotic device vertically. The way participants probed the force fields was also not restricted and they could choose the way they moved; for example, in a rhythmic or discrete manner.

The experiment was divided into two sessions of 180 trials, completed in two days. To become familiarized with the experimental setup and to ensure that the participants understood how to grasp the device, they performed a training set at the beginning of each day. The training set consisted of two repetitions of the ten levels of stiffness for the *comparison* force field, and a 0 mm/m tactor displacement gain for the *standard* force field. During training, we provided feedback to the participants about their responses. For the test trials, we had 40 different force fields pairs (10 *comparison* stiffness levels and 4 *standard* skin stretch gain levels) that were repeated eight times throughout the test session. The overall experiment consisted of 40 training trials that were not analyzed, and 320 test trials. All the different conditions were presented in a pseudorandom and predetermined order.

### Experiment 2

Ten participants (seven females, 22-26 years old) participated in the experiment. This experiment focused on the predictive and reactive components of grip force control due to the skin-stretch stimulus. To dissociate the contribution of the feedforward and feedback grip force adjustments, we incorporated stretch-catch probes where we maintained the load force but unanticipatedly omitted the skin-stretch (Hermsdörfer, Blankenfeld 2008). The protocol of this experiment was similar to the protocol of Experiment 1, with a few changes. We asked the participants to distinguish between pairs of force fields but we only used three stiffness values for the *comparison* force field (40, 85, and 130 N/m). For the *standard* force field, we used a constant stiffness value of 85 N/m with tactor displacement gains of 0, 33, 66, and 100 mm/m. We asked participants to perform eight discrete movements in each of the two force fields. We only counted movements that started and ended outside of the elastic field, and were completed in 300 msec and extended at least 20 mm into the force field. In addition, we presented a counter of successful movements to the user. After eight probing movements with the first force field, the screen automatically switched to the second force field. When participants had performed the eight probing movements into the second force field, the screen automatically became white and participants had to choose the force field they perceived as stiffer.

The experiment consisted of 132 trials, 24 training trials and 108 test trials, and was divided into two sessions of 66 trials, completed in two days. The protocol for the test study consisted of twelve pairs of force fields (three levels of stiffness for the *comparison* force field and four levels of tactor displacement gain for the *standard* force field). Each pair of *comparison* and *standard* fields was repeated in nine trials throughout the experiment. In six of the nine trials, we introduced stretch-catch probes either in the second or in the seventh movement, in which we maintained the load force but unanticipatedly omitted the skin-stretch. In three of the six trials, the stretch-catch probe was the second probing movement of the total of eight probes, and in the other three, it was the seventh probing movement. Overall, 3.125% of the probes were stretch-stretch-catch probes. We chose to introduce stretch-catch probes in the second and not the first probing movement since it would have been meaningless to omit the tactile stimulus before participants felt it at least once. All the different conditions were presented in a pseudorandom and predetermined order.

## Data Analysis

We used the Lilliefors test to determine whether the dependent variables were normally distributed (Lilliefors 1967), and found that all the dependent variables came from normal distributions.

### Experiment 1

#### Perception

For each participant, we fitted psychometric curves for the probability of responding that the comparison force field was stiffer as a function of the difference between the stiffness of the comparison and the standard force field using the Psignifit toolbox 2.5.6 (Wichmann, Hill 2001). We repeated this procedure for every level of tactor displacement gain. To assess the effect of the artificial skin-stretch on perception of stiffness, we computed the point of subjective equality (PSE) and the just noticeable difference (JND) of each psychometric curve. The PSE indicates the difference in stiffness levels at which the probability of responding that the comparison force field had a higher level of stiffness was 0.5. A positive PSE value; i.e., a rightward shift of the psychometric curve, represents overestimation of the standard force field, and a negative PSE value; i.e., a leftward shift indicates an underestimation. The JND was measured as half the difference in the comparison stiffness levels between the 0.75 and the 0.25 probabilities of responding that comparison force field had a higher level of stiffness. The JND indicates the sensitivity of the participant to small differences between the stiffness levels of the two force fields.

After extracting four PSE and four JND values for each participant (for each level of tactor displacement gain), we fitted regression lines to these values. To test the significance of the change in PSE and JND values due to tactor displacement gain, we used a repeated-measures regression with the *anovan* MATLAB function. The independent variables were tactor displacement gain (continuous), and participants (random). The model also included the interactions between the variables.

#### Action - Peak grip force–peak load force ratio

We filtered the recorded grip force data using the MATLAB function *filtfilt* with a 2^nd^ order Butterworth low-pass filter, with a cutoff frequency of 15Hz, resulting in a 4^th^ order filter, with a cutoff frequency of 12.03 Hz. On each trial, we examined the load force generated by the robotic device, the magnitude of tactor displacement, and the grip force that participants applied. The grip force trajectories had a non-uniform peaked pattern that appeared predominantly in trials in which skin-stretch was applied. Therefore, we used the MATLAB function *findpeaks* to identify the grip force and load force peaks and then manually corrected by visual examination. To evaluate the grip force modulation during the interaction with the *standard* force field, we calculated the peak grip force-peak load force ratio of the first and last probing movements for each level of tactor displacement gain. The participants could not predict the tactor displacement gain used for the *standard* force field in the first probing movement, whereas in the last probing movement, participants already had information about the stimulus and likely formed an internal representation of it. To test the significance of the change in the applied grip force due to tactor displacement gain and between probing movements, we used a repeated-measures ANCOVA using *anovan* MATLAB function. The independent variables were tactor displacement gain (continuous), the probing movement (first or last, categorical), and participants (random). The model also included the interactions between the independent variables.

### Experiment 2

In the next analyses, we separated the probing movements and analyzed the grip force trajectories associated with the applied load force. We identified the start and the end of each probing movement using the load force signal. The load force was equal to zero when the position of the device was greater than zero, and increased as a function of the penetration when the participants crossed the boundary of zero. The analyses were only run on data collected during the interactions with the *standard* force field that had a constant stiffness value.

#### Feedforward grip force control

To isolate the predictive component of grip force control due to the skin-stretch stimulus, in the first two analyses in this experiment, we only examined the applied grip force from probing movements with stretch-catch probes. For a given trial, we examined the load force applied during interactions with the force field, and the applied grip force. To test the change in grip force control between the initial and final probing movements of each interaction, we analyzed the second and seventh stretch-catch probing movements separately. Visual examination of the zero stretch trials suggested that there was a rapid change in grip force-load force ratio between the first and the second probing movements. However, because we could only incorporate stretch-catch probes starting from the second probe, in trials with skin-stretch we could not examine the grip force in the first probing movement. Therefore, in this experiment the analyses of trials with a skin-stretch stimulus (gains: 33, 66, and 100 mm/m) were examined independently of trials without skin-stretch (gain 0 mm/m). In trials with skin-stretch we compared the second to the seventh probing movements, and for reference, in trials without skin-stretch, we compared the first, second, and seventh probing movements.

To quantify the effect of skin-stretch during the repeated exposure to the elastic force field, we performed two analyses. To match the analysis of Experiment 1, we computed the *peak grip force-peak load force ratio.* The grip force is generally coupled with load force but may precede it; hence to ensure we had captured the beginning of the grip force change, we included 50 samples before and after the start and end movement points.

To isolate the modulation of grip force with load force from the baseline grip force control, we analyzed the *grip force-load force regression*, similar to the analysis in (Flanagan, Wing 1990, Leib, Karniel & Nisky 2015, Nowak et al. 2002). In this analysis, to be sure that we only examined the forces applied during the interaction with the force fields, we did not add 50 samples before and after the start and end points. For each participant, for the second and seventh probing movements on each of the 108 trials, we fitted a two degrees-of-freedom regression line (slope and intercept) to the trajectory in the grip force-load force plane. For the slope and intercept separately as a dependent variable, we fitted a repeated-measures ANCOVA model using *anovan* MATLAB function. The independent variables were tactor displacement gain (continuous), the probing movement (second or seventh, categorical), and participants (random). The model also included the interactions between the independent variables.

#### Feedback grip force control

To quantify the component of grip force control elicited in reaction to the stretch, we subtracted the grip force trajectories in the second probing movement with only load force (stretch-catch probe) from the grip force trajectories of the second probing movement with both skin-stretch and load force stimuli. For each tactor displacement gain, we considered all the second probing movements without stretch-catch probes and all the second probing movement with stretch-catch probes separately. We also subtracted from each trajectory the grip force at the beginning of the normalized trajectory. Then we subtracted the average grip force trajectory as a result of the load force stimulus from the average grip force trajectory as a result of skin-stretch and load force stimuli, such that the remaining signal was the reactive effect of the stretch stimulus on grip force.

Because of the close proximity between the tactor displacement and the measurement of grip force, we suspected that there was an artifact of the tactor movement in our grip force measurements. Therefore, both the artifact and the reactive response of the user could have affected the calculation of the reactive response. Accordingly, we characterized the artifact of our measurement system to movement of the tactor (see SI), and subtracted it from the reactive component.

To quantify the feedback response of the users to stretch stimulation for different values of tactor displacement gain, we calculated the difference between the reactive grip force at the end of the interaction with the elastic force field and the minimum of the grip force trajectory. This measure allowed us to quantify the entire range of change in grip force in response to tactor displacement, which we then termed the *reactive grip force range*. To test the dependence of the *reactive grip force range* on the different levels of tactor displacement gain, we fitted a repeated-measures regression model using *anovan* MATLAB function. The independent variables were tactor displacement gain (continuous) and the participants (random). The model also included the interactions between the independent variables.

## Supporting Information

### Artifact characterization experiment

To quantify the tactor displacement artifact on grip force measurement without human contact, we conducted a validation test in which we applied a constant grip force of 1N, using a padded clamp that resembled the finger-pad grip, and measured the grip force during tactor movement. We held the handle of the haptic device well above the skin stretch device, and performed a number of interactions with the force field that had a constant level of stiffness of 85 N/m, for each tactor displacement gain level. The purpose of this manual interaction was to make sure that we were using a human-like probing trajectory similar to the ones that the participants employed during the experiment.

To characterize the artifact response during the interaction with a force field and without human contact, we analyzed the grip force trajectories associated with the applied load force, similar to our analysis of grip force trajectories in the experiment. Both the load force and tactor displacement changed as function of the penetration into the force field. Hence, to compare the artifact for different levels of tactor displacement gain we normalized the grip force signal by dividing the measured grip force by the peak load force of each probing movement. In addition, the trajectories were time-normalized and aligned such that 0 was the onset of the contact with the force field, and 1 was the end of the interaction. This resulted in artifact measurements that were expressed in the same units as the grip force measurements during the experiment.

We identified the start and the end of the probing movement using the load force trajectory. The load force was equal to zero when the position of the device was greater than zero, and increased as a function of the penetration when the participants crossed the boundary of zero. Similar to our analysis of the grip trajectories in the regular experiments, we included 50 samples before and after the start and end movement points.

Fig. SI1 presents the average grip force artifact trajectories normalized by the peak load force. The average grip force trajectories had a constant pattern: when the tactors started moving, there was a steep decrease in the measured grip force followed by its recovery and a moderate increase. The size of the artifact in the measured grip force depended on the tactor displacement gain; thus, in our analysis of the reactive grip force components, we subtracted the characterized artifact for each tactor displacement gain.

**Fig. SI1.**
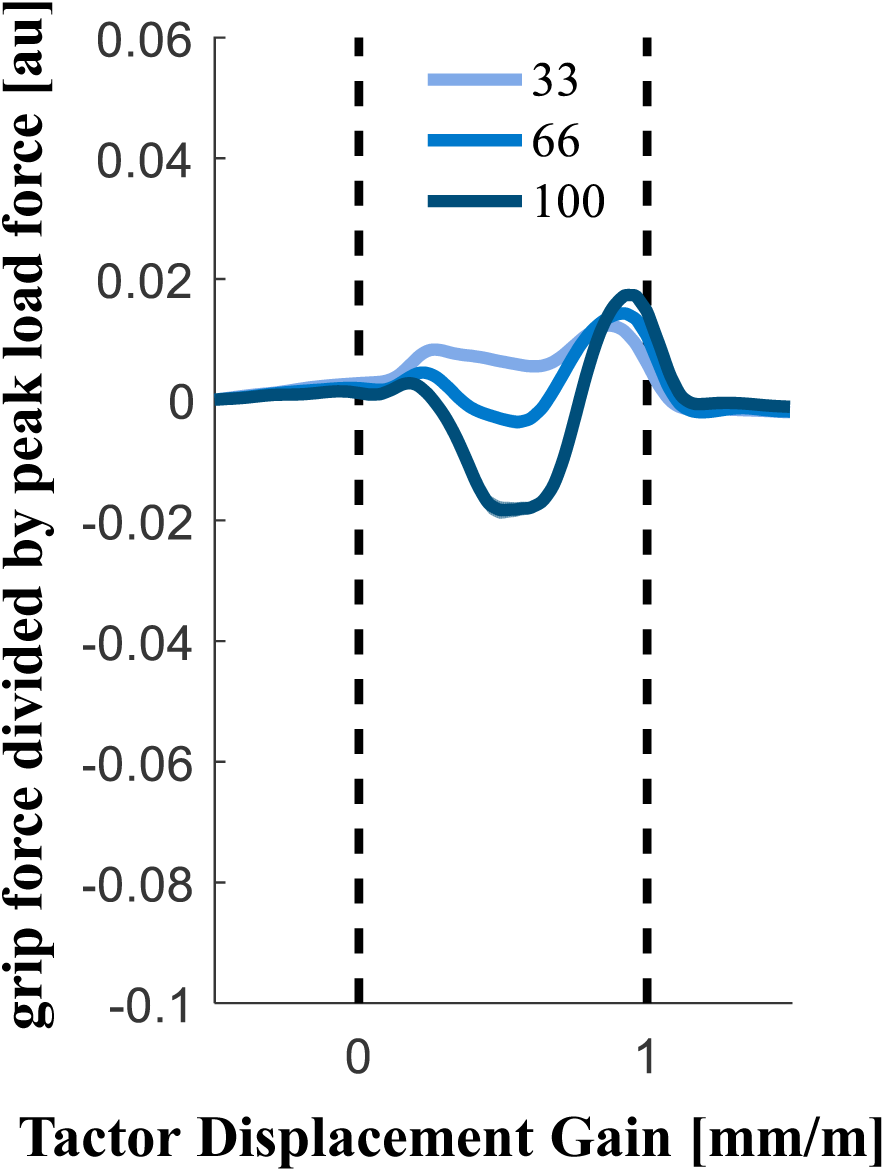
The artifact of tactor movement on measured grip force without human contact. Average grip force trajectories divided by peak load force. Shading represents the standard errors, and the black vertical dashed lines represent the time of the interaction with the elastic force field.

## Acknowledgments

The authors would like to thank Amit Milstein, Zhan Fan Quek, and Eli Peretz for their help and advise in designing and building the skin stretch device, and Guy Avraham for valuable comments on the manuscript. The study is supported by the Binational United-States Israel Science Foundation (grant no. 2016850), by the National Science Foundation (grant no. 1632259), by the Israeli Science Foundation (grant 823/15), and by the Helmsley Charitable Trust through the Agricultural, Biological and Cognitive Robotics Initiative of Ben-Gurion University of Negev, Israel. Mor Farajian was supported by the Tzin Fellowship.

